# Downscaling local distribution of cattle over Guadeloupe archipelago: an adapted method for disaggregating census data

**DOI:** 10.1101/2025.05.02.651856

**Authors:** Dufleit Victor, Guerrini Laure, Marius Gilbert, Daniele Da Re, Etter Eric

## Abstract

Gridded livestock distribution datasets have been produced for several years and are used in various fields, including epidemiology, livestock impact assessment, and territory management. Those datasets are based on census conducted at national/sub-national scale which are then downscaled using machine learning algorithm and relevant spatial-explicit environmental predictors. The most known dataset of livestock disaggregated observations is the Gridded Livestock of the World (GLW), which produces global maps of livestock density at 10 km spatial resolution for several livestock species. Though this spatial resolution can be appropriate to describe livestock distribution at the global scale, it inherently leads to a coarse representation of breeding species density for smaller territories such as the Caribbean Islands.

In this study, we propose an adaptation of the GLW methodology that accounts for the spatial autocorrelation in observed cattle distribution, thereby better capturing the specific characteristics of geographically limited areas such as the Guadeloupean archipelago. Cattle census data were collected for the 32 municipalities of the archipelago and associated to environmental predictors derived from remote sensing and land cover datasets. Together with the Random Forest (RF) algorithm used in the standard GLW methodology, we tested the performance of a Geographical Random Forest (GRF), a novel methodology allowing for taking into account the spatial autocorrelation of the response variable. The GRF algorithm demonstrated significantly better performance compared to the RF algorithm, albeit with longer processing times, and allowed us producing cattle distribution maps for the entire Guadeloupe archipelago at a spatial resolution of 225 m using both algorithms. The approach developed holds potential for application to other small territories, including other islands in the Caribbean.

## Introduction

Knowing the spatial distribution of livestock abundance is a key information to guide and implement livestock production and sanitary policies. National or local administrations conduct livestock censuses to obtain detailed information on animal densities and distribution, which are then used to assess the development and the economic benefits of the livestock sector, to measure the impact on natural resources or on public health and to control livestock diseases (1).

The GLW database, first released in 2007, provides a gridded and downscaled representation of livestock densities worldwide (2–4). Livestock density estimates in GLW are derived from census data collected at different scales (national or sub-national - region, district, communal scale/village) and downscaled using environmental covariates and statistical methods. The first version of the GLW (2) applied a stratified regression approach to estimate livestock densities at a spatial resolution of approximately 5 x 5 km at the equator (3 minutes of arc). In contrast, newer versions of GLW (4, 6) utilize the Random Forest (RF) algorithms to downscale livestock census globally at spatial resolution of approximately 10 × 10 km at the equator.

While this spatial resolution choice reduces spatial artifacts in predictions at a global scale, it results in a coarse representation of livestock density in smaller geographical areas like the Caribbean islands, where detailed livestock density information is critical for risk assessment and sanitary policies (5). For example, in Guadeloupe, an overseas department and region of France in the Caribbean, the GLW dataset represents the entire archipelago with just 15 pixels.

Another limitation of the current GLW is its inability to explicitly account for spatial autocorrelation of the response variable. The RF algorithm does not inherently account for spatial autocorrelation. Though spatial proxies such as coordinates of training observations or distance between observations could be added as predictors to account for potential spatial autocorrelation in the training dataset, this approach has not always lead to gain in prediction accuracy or pertinent spatial pattern reproduction (6). Various model formulation have been proposed to incorporate spatial information into RF (7,8), among which the Geographical Random Forest approach (GRF) developed by Georganos et al. (2021) (9) stems out. GRF addresses the limitation of standard RF by integrating spatial interactions between variables, thereby improving prediction accuracy.

Despite these advances, the use of such models in data-scarce and spatially heterogeneous contexts—such as small island territories—remains limited. This is particularly relevant for regions like the Guadeloupe archipelago, where traditional livestock data are often fragmented or unavailable at high spatial resolution. In such settings, evaluating the performance of models like RF and GRF becomes crucial for developing reliable, context-sensitive predictive tools.

In this study, we assess the applicability of predictive models trained on small datasets and propose a methodological strategy to mitigate issues related to spatial autocorrelation. Specifically, our goal was to develop a Territorial Livestock Mapping (TLM) model for the Guadeloupe archipelago using both RF and GRF algorithms, to predict cattle densities at a fine spatial resolution of 225 meters. These high-resolution maps aim to support epidemiological research by offering a detailed spatial representation of livestock distribution across the territory. In addition, we evaluated the impact of methodological choices on the predictive performance of the two algorithms. Specifically, we tested the impact of the choice of i) different sampling strategies and ii) various input variables, including topographic descriptors, land cover data, and environmental parameters, selected based on their ecological relevance to cattle distribution and habitat suitability.

## Materials and Methods

### Study Area

The archipelago of Guadeloupe is located in the Caribbean and is part of the French West Indies. It comprises “mainland” Guadeloupe, with two islands, Basse Terre (848 km², to the west) and Grande Terre (588 km², to the east), plus the administrative dependencies of the islands of La Désirade (21 km²), Marie-Galante (158 km²) and Les Saintes (14 km²). The tropical climate is influenced by the sea and trade winds with microclimatic variability driven by topography. Basse-Terre is dominated by the volcanic massif of La Soufrière (1467 m), resulting in a rainy climate, with annual rainfall ranging from 1000 to over 7000 mm, picking at the volcano’s summit. In contrast, Grande Terre’s lack of significant relief, results in a drier climate with annual rainfall between 1000 and 1500 mm(14). The administrative dependencies, share a similar climate to Grande Terre. Guadeloupe has a population of around 400,000 with tourism as its primary economic activity. Agriculture remains important, with two main crops, sugar cane and bananas (15). Livestock farming has decline since the 1980s (16) due to factors like urban development, tropical parasites such as ticks (e.g. *Amblyomma variegatum, Rhipicephalus microplus* (18)) and related disease (e.g. *anaplasmosis, heartwater*… (19,20)), feral dog attacks (17) and livestock theft (18). While tethered rearing dominates, grazing is also practiced. Traditionally tied to sugar cane cultivation, cattle are now used for subsistence and leisure activities, with the tradition of “ox pulling”. These challenges, combined with land access issues, have contributed to the decline of cattle farming in Guadeloupe.

### Census data

The cattle census data, extracted from the database collected by the departmental livestock institute (2021 census; Agriculture, Alimentation and Forest Direction (DAAF), and obtained from DAAF Guadeloupe in 2023, provided a list of 5 320 cattle breeders, representing a total of 37 139 cattle heads. Data without information on the location of the breeders (i.e., municipalities) were not included in the survey. Data included the identification of the breeder (EDE number), its location (municipality name) and the number of cattle per breeder. Cattle density per municipality was calculated (Figure 1). Administrative boundaries were obtained from BD TOPO® (19). Moran Index was computed to account for potential spatial autocorrelation in the dataset (20).

**Figure 1.**
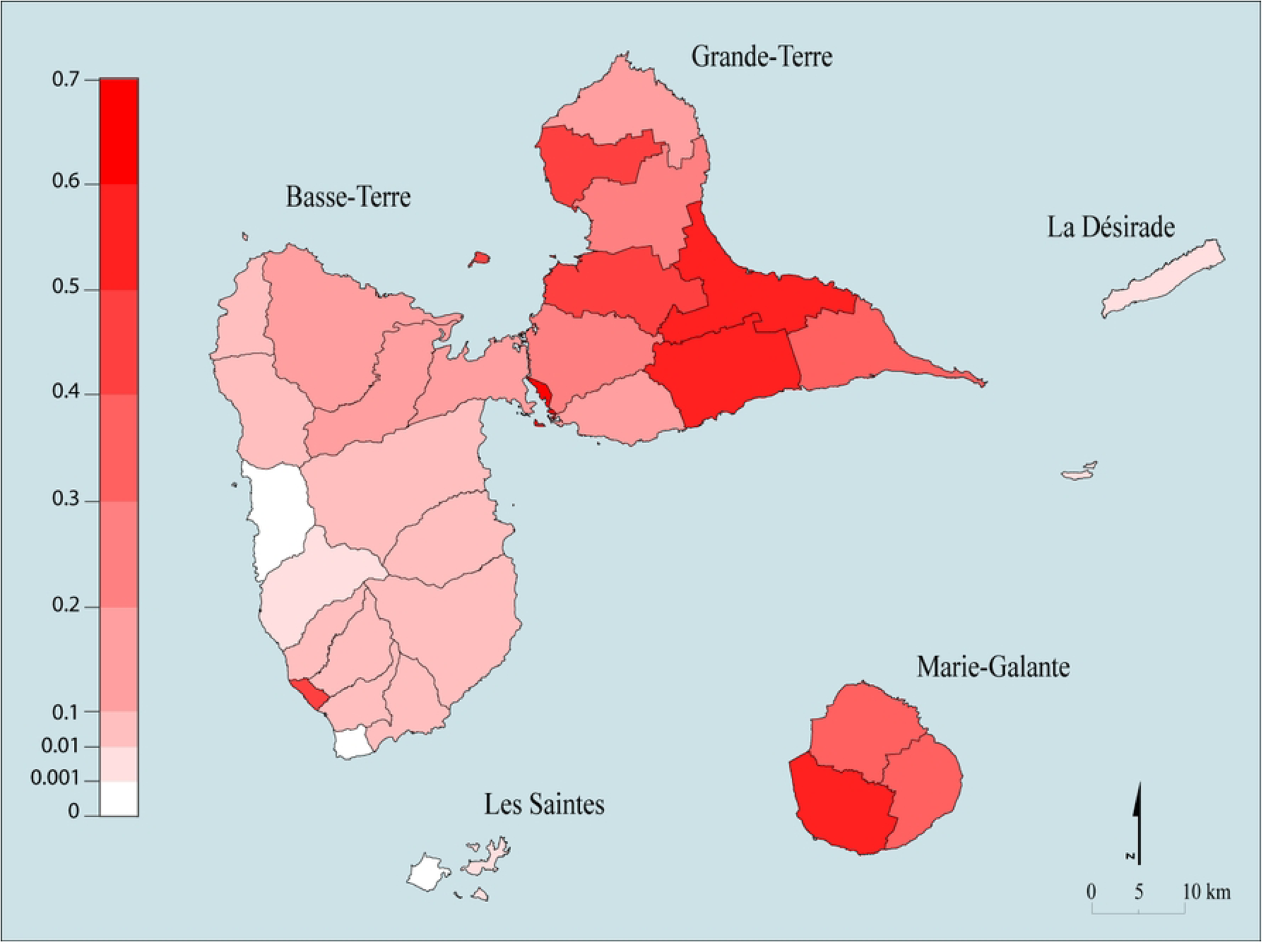
Observed cattle densities in Guadeloupe municipalities.

### Suitability raster & Masks

Masks are commonly employed in studies or modelling processes to restrict the area of interest and the area of training and extrapolation of the model. In this study, a mask was created to delineate areas within the Guadeloupean territory where cattle breeding was either feasible or not. The construction of this mask was guided by the methodology outlined below:

- Land cover classification: The original land cover classes from the Karucover dataset were reclassified into two categories: “Suitable” or “Unsuitable” for cattle breeding (S1). Land cover classes were analysed to determine the suitability of natural vegetated areas, such as grasslands and forests. Land use data were incorporated to improved classification accuracy (S1a). Agricultural land and low-density home gardens were classified as suitable, as cattle are occasionally raised on fallow agricultural land or in-home gardens. Specific vegetated public infrastructures (e.g., roadside vegetation or roundabouts) were also considered suitable for grazing, particularly for farmers with limited herds or land availability (S1b).
- Exclusion of unsuitable areas: Areas where cattle breeding is unfeasible, such as the National Park and urban zones (19,20; S2), were excluded from the analysis (S1c ; S2).
- Suitability raster creation: A raster with a resolution of 225 x 225 m was created to cover the entire study area. Pixels were assigned a value of 1 (suitable) if they contained at least one suitable polygon (S1d). This raster is referred to as the suitability raster (S2).

### Predictors

Spatially explicit predictors are crucial for generating cattle density prediction maps through downscaling approaches (22) (Table 1).

**Table 1.** Predictor variables used in the model. ,the “Predictor dataset” column shows the inclusion of different variables in the models (model I and model II) (see Experimental design part)

| Variable Name |  | Type | Use | Source | Predictor Dataset |
| --- | --- | --- | --- | --- | --- |
| Digital Elevation Model | DEM | Topographic | Spatial predictor | BRGM (25) | I ; II |
| Slope |  | Topographic | Spatial predictor | Created from DEM | I ; II |
| Topographical Ruggedness Index | TRI | Topographic | Spatial predictor | Created from DEM | I ; II |
| Ten land cover classes derived from Karucover 2017 |  | Land cover | Spatial predictor | Karugéo (13) | I |
| Barn distance raster |  | Barn distance raster | Spatial predictor | Karugéo (13) | I |
| Ponds Distance raster |  | Ponds Distance raster | Spatial predictor | Karugéo (13) | I |
| Road Distance raster |  | Road Distance raster | Spatial predictor | BD TOPO® (19) | I ; II |
| River distance raster |  | River distance raster | Spatial predictor | BD TOPO® (19) | I ; II |
| 10 Fourier-derived variables from Normalized Difference Vegetation Index | NDVI | Vegetation | Spatial predictor | MODIS (26) | I ; II |
| 10 Fourier-derived variables from Enhanced Vegetation Index | EVI | Vegetation | Spatial predictor | MODIS (26) | I ; II |
| 10 Fourier-derived variables from Day Land Surface Temperature | DLST | Thermal | Spatial predictor | MODIS (27) | I ; II |
| 10 Fourier-derived variables from Night Land Surface Temperature | NLST | Thermal | Spatial predictor | MODIS (27) | I ; II |

Topographic predictors were derived from a Digital Elevation Model (DEM) for Guadeloupe, provided by the geological and mining research bureau (Bureau de Recherche Géologique et Minière; BRGM). This included elevation, slope, and the Topographical Ruggedness Index (TRI), potentially influencing cattle distribution, by determining accessibility, ease of movement, and suitability for grazing. Steeper slopes or rugged terrain may restrict cattle grazing activities.

Land cover data from the high-resolution Karucover 2017 dataset (13), which includes 24 land cover classes and 53 land use classes, provides essential information about vegetation and land use types critical to cattle. From this dataset, ten land cover classes relevant to cattle breeding were identified in consultation with livestock specialists and used to generate 225 m resolution raster layers (see Tables S3, S4 and S5 for class conversions).

Distance rasters for barns, ponds (extracted from the Karucover 2017 dataset) as well as roads and rivers (extracted from the BD TOPO® dataset (16)), influencing cattle accessibility and breeding feasibility, were included.

Environmental variables were extracted from Moderate-resolution Imaging Spectroradiometer (MODIS) data (25), covering five years (2015–2019). Day/Night Land Surface Temperature (DLST, NLST) and vegetation indices such as the Normalized Difference Vegetation Index (NDVI) and the Enhanced Vegetation Index (EVI), capture seasonal dynamics in vegetation growth and thermal conditions. Temporal Fourier Analysis (TFA) of these indices allows the integration of ecological dynamics such as the annual growth cycle of crops (e.g., sugarcane fields) that impact cattle feeding areas. Therefore the MODIS time series were resampled to a 225 m resolution when necessary and summarised using TFA (23). This process enabled the calculation of metrics such as the mean, annual minimum and maximum, standard deviation as well as amplitude and phase of the annual, biannual and triannual cycles for the four selected indicators, which were then included in the modelling procedure (24).

A total of 57 predictor rasters were created for the entire archipelago of Guadeloupe.

A correlation matrix of the predictors and correlation between predictors and cattle densities at municipality level were calculated, based on a non-parametric measure of rank correlation (Spearman’s ρ).

### Disaggregating census Data: Experimental design

The methodology used was adapted from the GLW3 model (28) to a smaller scale to develop a Territorial Livestock Mapping model for the Guadeloupe islands. A summary of the method is presented in Figure 2.

**Fig 2.**
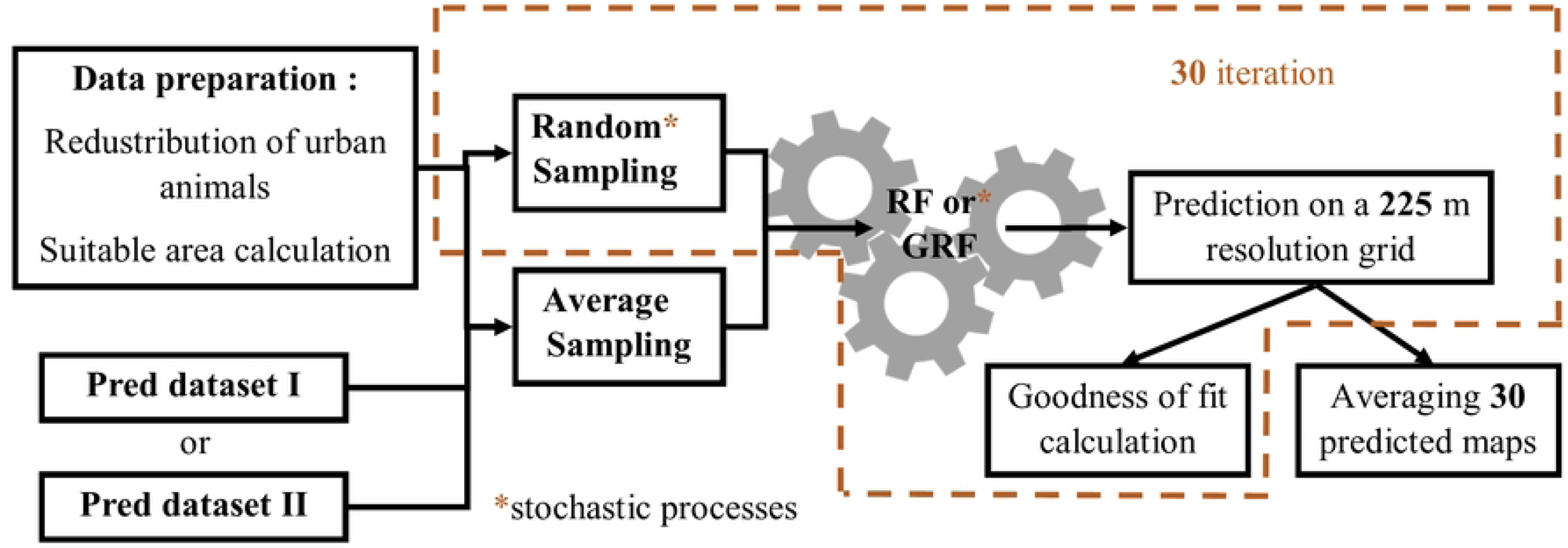
Flowchart of the study.

Preliminary exploration of census data revealed that the highest cattle densities were recorded in urban areas of Basse-Terre and Pointe-à-Pitre, reflecting the practice of breeders residing in urban centres while maintaining livestock in the surrounding rural areas. To mitigate potential artefacts, cattle were redistributed from urban municipalities to neighbouring rural areas using the following equation (Eq. 1):

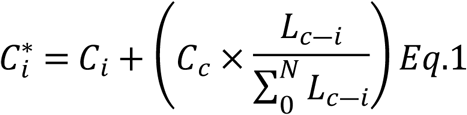

Where *C*_*i*_is the number of cattle in the *i*^th^ neighbouring municipality, *C_c_* is the number of cattle in the *c*^th^ urban municipality, N is the total number of neighbouring municipalities of an urban municipality, and *L*_*c*―*i*_is the length of the border between the *c*^th^ urban municipality and the *i*^th^ neighbouring municipality. Using the redistributed data and suitable areas (calculated as the total area of all suitable pixels per municipality based on the suitability mask), cattle densities were estimated for suitable areas. These calculations formed part of the data preparation process illustrated in Figure 2. Moran index was adjusted with the obtained ‘adjusted’ municipal cattle densities.

#### Sampling Predictor

We employed to two predictors sampling approaches (Figure 3) in order to assess the effect of this methodological choice on the models’ prediction accuracy (Da Re et al., 2020)(28).

**Fig 3.**
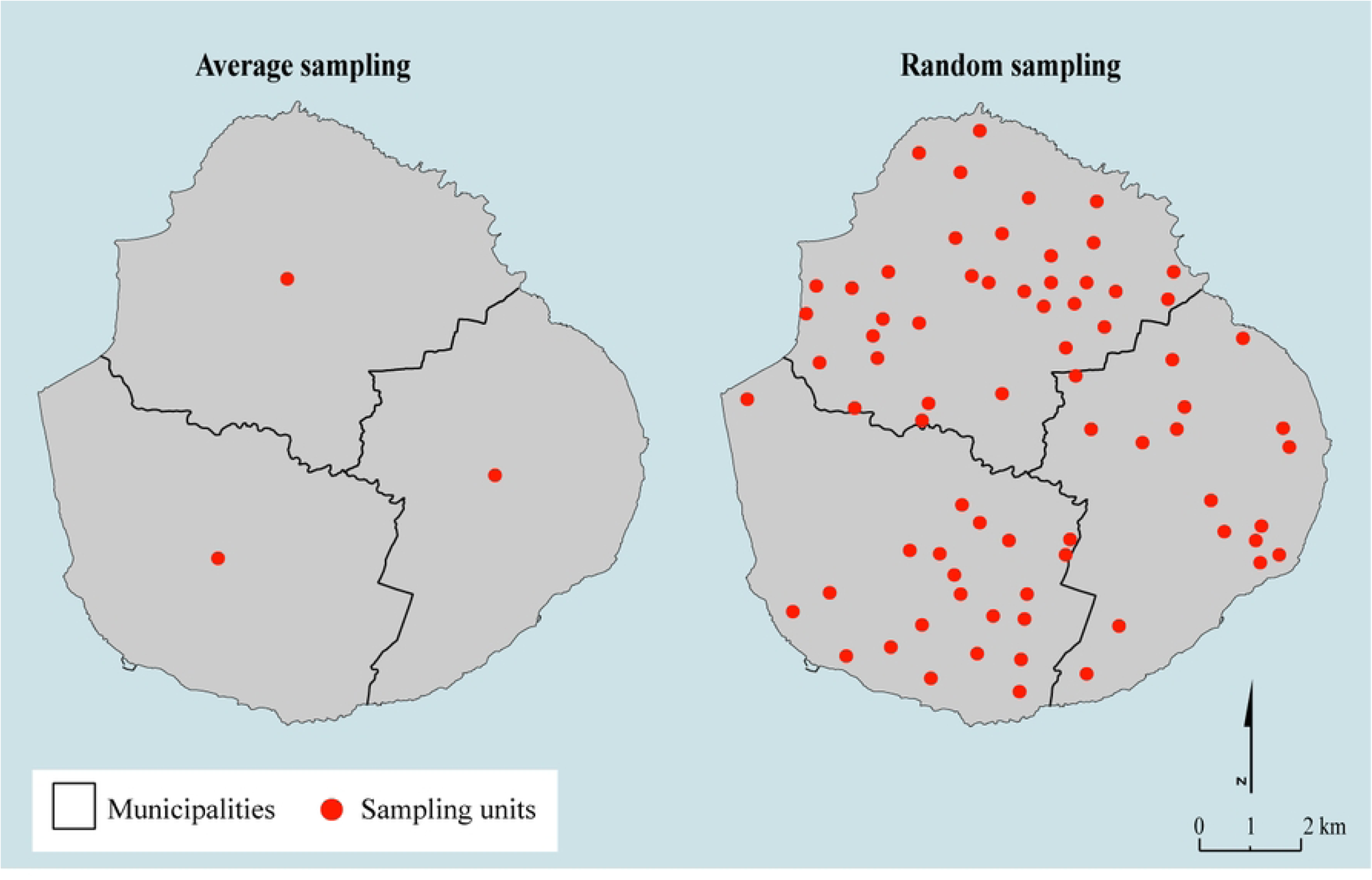
Presentation of the two predictors sampling methodologies.

The first approach, the average sampling method, involves calculating the average of predictor values over all the pixels within the polygon. This average sampling approach was previously used in studies by Li et al. (2021) and Da Re et al (2020) (11,28). The second approach, the random sampling method instead involves generating a randomly distributed number of points in the area of interest, in our case with a density of 50 points per 10 000 ha. Each point was associated with the predictor values derived from the pixel where it was located. The latter approach increased the size and variability of the training dataset. Random sampling was repeated 30 times to assess variability, while average sampling was conducted once, given the lack of stochasticity inherent to the latter process. For each dataset obtained through random sampling, one RF/GRF was trained, resulting in a total of 30 models. Using the average sampling dataset, 30 random forests were generated.

#### Modelling

The training sample, independently from the sampling strategy and predictors choice, was provided to one of the selected RF algorithms. The ranger R package (29) was selected for the standard RF model, while R package spatialML R package (31) was selected for the Geographical Random Forest (30) was. A standard RF model requires The hyperparameters of the standard RF were set as follows (Nicolas et al. (22).):

◦ **Number of trees (ntree)**: this parameter determines the number of trees to be grown. It was set as the number of training points divided by 20, with a minimum of 100 trees.
◦ **Node size**: the node size parameter influences the minimum number of samples required at each node after a split during tree construction. If after a split, a node has a size (number of samples) less than the node size, no further splits will be performed on that subsample. The node size was set to the number of samples divided by 1000 with a minimum node size of 3.
◦ **mtry**: the mtry parameter defines the number of predictors used to build each tree. mtry was set as the number of predictors divided by 3 with a minimum of 3 predictors.

The GRF model incorporated a “bandwidth” parameter, which defines the maximum distance between a data point and its neighbouring observations. Local models were created for each training sample location using an “adaptive” kernel (30). For each local model, points were collected around the locations using the K nearest neighbour rules. The “grf.bw” function from the spatialML R package was used to optimise the bandwidth values with a minimum value of 1/3 and a maximum value of 2/3 of the training data points (only five steps were tested to optimize the bandwidth). For each bandwidth tested, the coefficient of determination (r^2^) was calculated and the bandwidth with the highest r² value was selected for the final model run. For the GRF model predictions, the weight given to the global and local models for the predicted value had to be chosen. As recommended by Georganos et al. (9), the weights were set to 0.75 and 0.25 for the global and the local models respectively.

For both algorithms, we decided to test two model formulations using two predictors’ datasets:

- Model I including all the 57 predictors
- Model II with a reduced set of 45 predictors, excluding Karucover derived predictors

The second model formulation was used to assess the accuracy of the prediction without high-resolution land cover data. This ensures the applicability of the model to other Caribbean islands with limited land cover information.

#### Validation

A predicted map, with a resolution of 225 m was generated at the end of all bootstraps. To evaluate the relative importance of predictors, Permutation variable importance (PVI) was calculated during model training (32). While PVI highlights the relative contribution of each predictor to model accuracy, it does not indicate whether the effects on the dependent variable are positive or negative. To complement this analysis, PVI was compared with the Spearman correlation (absolute value) between observed municipal cattle density and predictors (averaged at the municipal level), calculated prior to modelling. Additionally, correlations between predictors were calculated using the full raster dataset. Predicted density maps were converted to animal count maps by multiplying the predicted cattle density by the pixel area. These maps were then aggregated at the municipality level to derive the total predicted animal counts per municipality. To evaluate model performance, the Roots Mean Squared Error (RMSE) and the Pearson r Correlation Coefficient (PCC) were calculated between observed and predicted municipal counts. RMSE assessed prediction accuracy (i.e., how closely predictions matched observed values), while PCC measured the proportionality between the observed and predicted values.

To compare the performance of GRF and RF, the distributions of RMSE and PCC were analysed across the eight combinations of modelling procedures. One-factor ANOVA and the associated Tuckey honestly significant difference (HSD) were used to determine statistically significant differences between methods. Observed and predicted values were standardised to the interval [0;1] using min-max standardisation (33), and correlation plots were then generated to compare outputs of the different methods.

#### Post-processing

The means and standard deviations of all the predicted maps were calculated to produce a final raster. This final raster served as a weighting layer to redistribute census data across the territory. This post-processing procedure, inspired by the GLW methodology, adjusted the predicted animal counts to ensure that the sum of all pixel values equalled the total number of animals in the Guadeloupe Archipelago. The adjusted pixel value *X_i_* was calculated using the following equation (Eq.2):

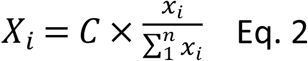

Where *C* represents the total animal count from the DAAF Census, n is the total number of pixels in the study area and *x_i_* is the predicted animal count for the *i*^th^ pixel.

The entire methodology was implemented using R © software version 4.2.3 (34).

## Results

### Produced maps and predictor importance

To run the models, 4 971 breeders, representing 34 790 cattle were finally used. A Moran Index of 0.61 (p-value < 0.05) was found using raw cattle densities data. Results of data preparations conducted before modelling (city animal redistribution and corrected density calculation) are displayed in supplementary materials (S6). Corrected cattle densities gave a Moran Index of 0.51 (p-value < 0.05). The predicted maps generated by the GRF model are shown in Figure 4 (RF results are not displayed as they exhibit similar pattern to those of GRF). The simulated animal count per pixel ranged from 0 to 2.93 using the random sampling method, and from 0 to 2.5 using the average sampling method.

**Figure 4:**
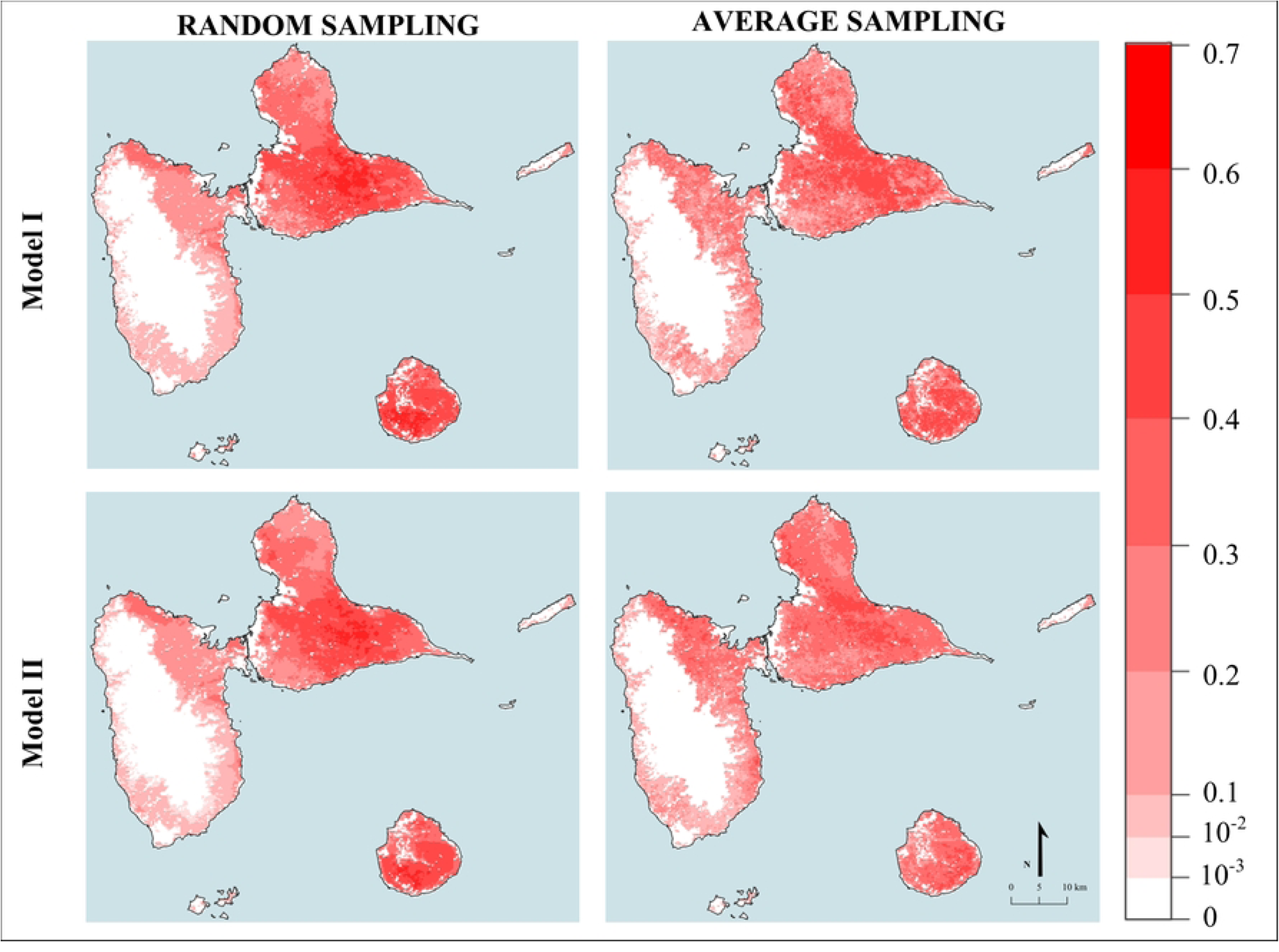
Predicted maps produced with GRF using two sampling methods and two predictors dataset.

For both sampling methods, the models effectively reproduced the observed distribution of cattle. The highest densities were observed on Marie Galante and in the central region of Grande Terre. These patterns correspond to the municipal densities calculated after redistributing animals from urban areas.

The Spearman correlations between cattle densities and the predictors, as well as the correlations between the predictors, are shown in supplementary materials (S7). The PVI is summarised in Figure 5. Significant positive correlations (p < 0.05) were found between cattle density and the breeding area predictor, which had the greatest impact on model I performance for both the RF and GRF algorithms. Significant positive correlations were also observed with agricultural land cover, which did not appear to be an important predictor in either model predictions. Distance to barn distance to pond, daily temperature standard deviation (DLST SD), slope and TRI were initially significantly negatively correlated with cattle densities (p < 0.05). The predictor Distance to barn was important in Model I particularly for the average sampling method. DLST SD proved to be an important predictor when applying the random sampling method particularly with model II. Slope appeared to be an important predictor in Model II whatever the sampling procedure. Distance to pond and TRI did not appear to be important predictors regardless of modelling method. Notably, grass cover, distance to river and mean night temperature showed particular relevance (Fig. 5) in the random sampling method, although no significant correlations with cattle density were initially detected.

**Fig 5.**
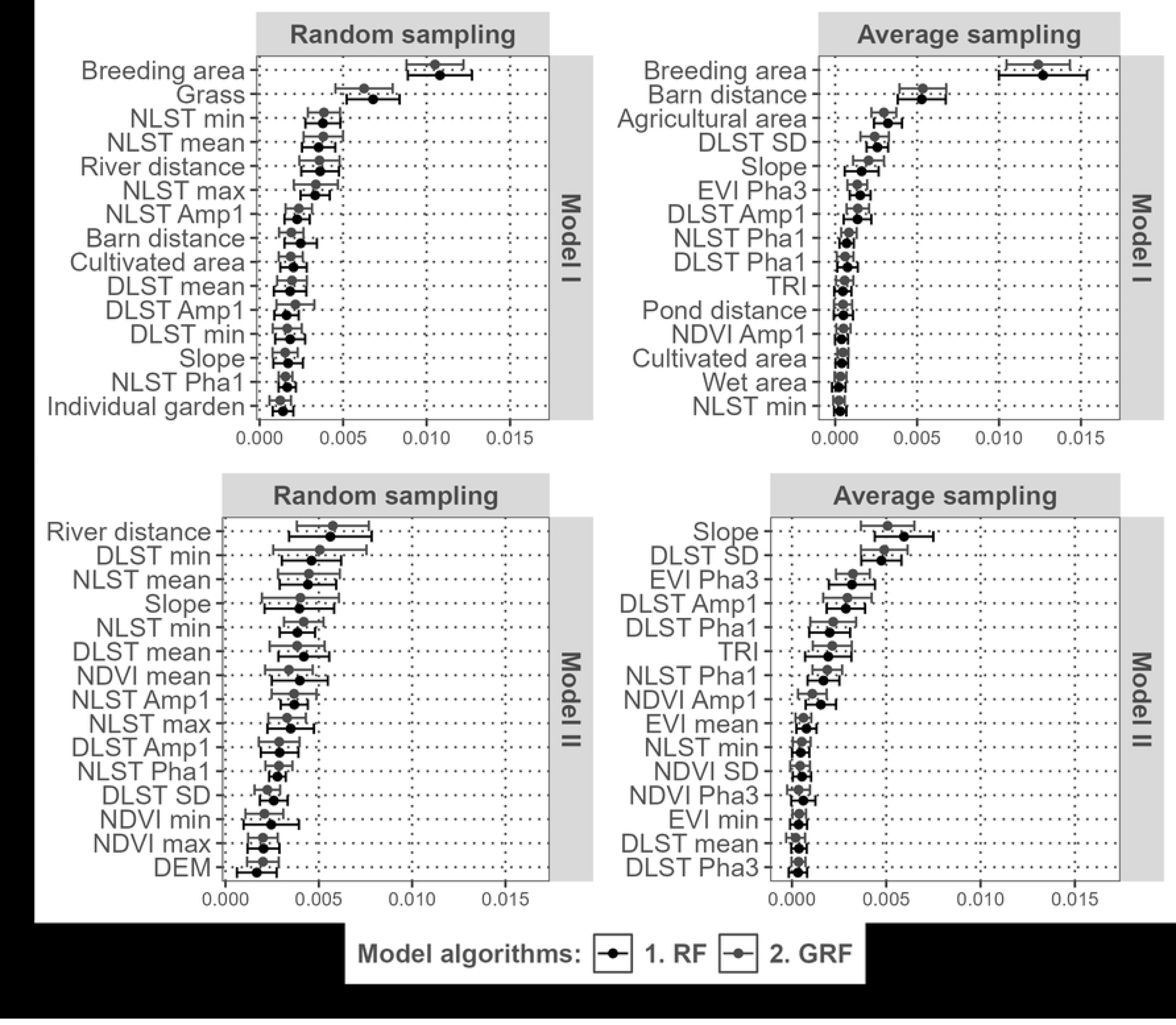
Permutation variable importance calculated by RF and GRF models. Only the 15 most representative variables are presented here. DLST = Day Land Surface Temperature; NLST = Night Land Surface Temperature; SD = standard deviation; Amp1= Amplitude of the annual harmonic; Amp2= Amplitude of the biannual harmonic; Amp3 = Amplitude of the triannual harmonic; Pha1 = Phase of the annual harmonic; Pha2 = Phase of the biannual harmonic; Pha3 = Phase of the triannual harmonic (16).

### Goodness of Fit measures (GOF)

The RMSE and PCC metrics produced consistent results across the tested methodologies. Higher Figure 6 shows that higher PCC values mean lower RMSE values, suggesting the model reliably followed the observed values. Both sampling methods resulted in high Pearson’s r values (> 0,9). Among the GOF metrics, the random sampling method demonstrated superior predictive performance, regardless of the random forest type (GRF and RF) or the set of predictors used (model I or model II). Within both average and random sampling methods, the GRF model provided more accurate predictions of cattle density compared to the RF model.

**Fig 6.**
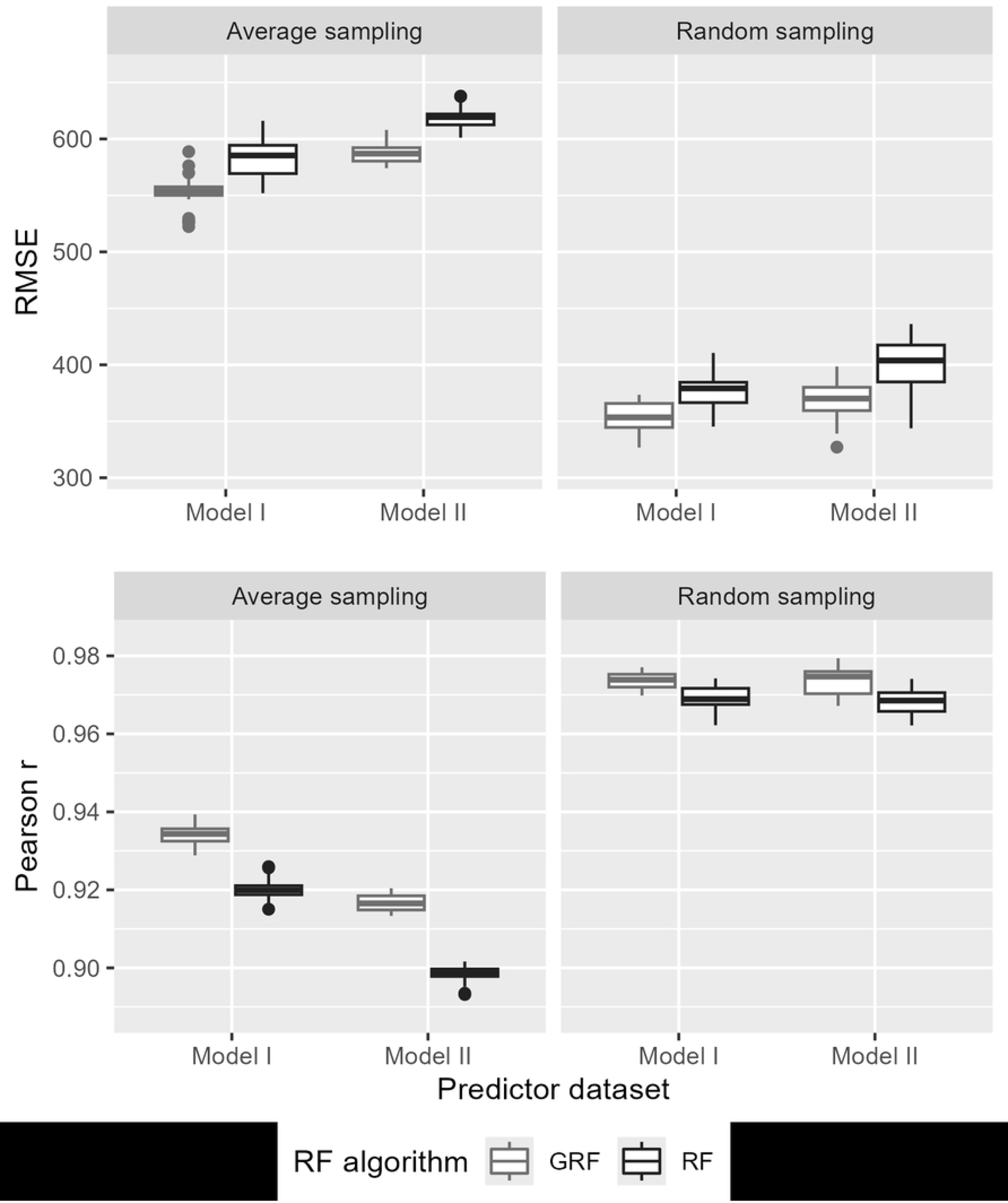
Synthesis of RMSE and Pearson correlation for explored methodologies. Different letters indicate statistically significant differences in the means as determined by one-way ANOVA and Tuckey HSD.

The inclusion of land cover predictors into the model (Model I vs. Model II) did not improve the GOF metrics when the random sampling method was used. Conversely, when the average sampling method was employed, Model I showed statistically higher PCC values than Model II.

Figure 7 compares the standardised simulated animal counts with observed counts at the municipal level. For both models, the two random forest methods combined with the two sampling methods, overestimated animal counts in municipalities with the lowest observed animals and underestimated it in the municipalities with the highest observed counts. The random sampling method provided the best fit, better simulating higher animal counts in municipalities with the largest observed population.

**Fig 7.**
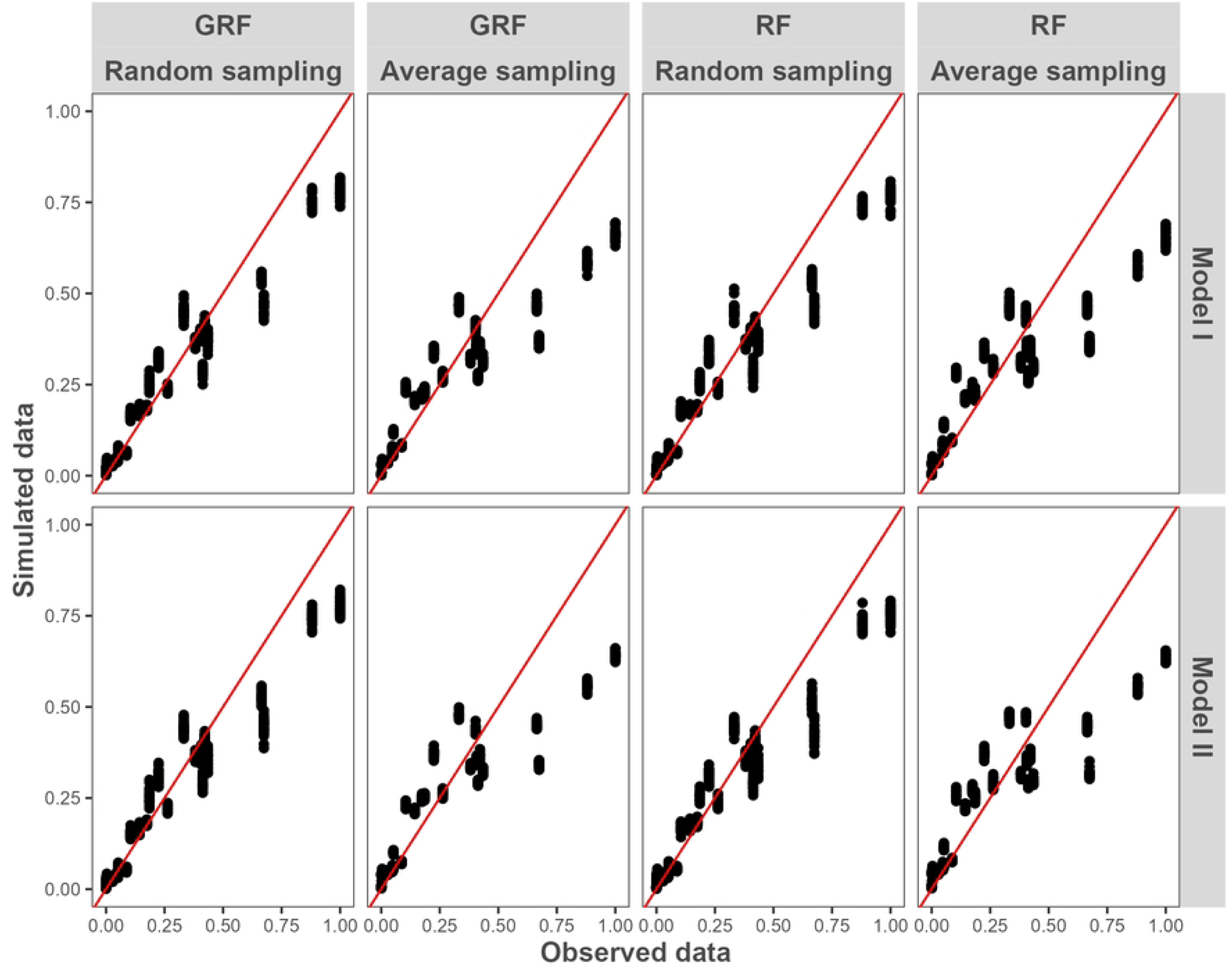
Standardized observed and simulated municipal cattle counts. The red line represents the equation f(x) = x

## Discussion

The RF and GRF census disaggregation methods were successfully applied to the Guadeloupean territory, resulting in the first cattle density map with a spatial resolution of 225 meters. This work represents a significant advance in livestock census disaggregation, as such methods had never before been implemented on such a small scale as evidence by the case study of the Guadeloupean archipelago.

### RF vs GRF

The GRF significantly outperformed classical RF in terms of predictive accuracy, regardless of datasets of predictors used and whether average or random sampling was used. Similar improvements in predictive performance with GRF over RF have been reported in different research areas (35), but with an increase in computational time (36). This highlights the critical role of spatial heterogeneity in improving predictive accuracy for population density modelling.

In fact, since spatial autocorrelation was observed in our census dataset (with Moran I = 0.61 after city animal redistribution (S6B) and a Moran I = 0.51 after calculation of density with suitable pixels (S6D), both p value <0.05), the inclusion of spatial proxies for our case study resulted relevant(7,8).

### Average sampling vs Random sampling

We also assessed the effect of the choice of two methods for sampling predictors on the random forest models predictive accuracy. Random sampling showed better GOF metrics and produced a wider range of predicted density values (0 to 2.92 animal per pixel compared to 0 to 2.46 with average sampling). This improvement may be attributed to the higher predicted animal counts in the most densely populated municipalities (Fig. 7). However, no major differences were observed for less densely populated municipalities. The better performance of the random sampling method could also reflect that this method might better took into account the spatial heterogeneity within municipality compared with the average sampling.

The model output maps showed different density patterns between the two sampling methods. This may be due to differences in the ranking of predictors according to the variable permutation importance when using different sampling methods. With average sampling, predictors previously identified as significantly correlated with cattle density (S7) also had high importance in RF models such as breeding area. Conversely, random sampling emphasised the importance of alternative predictors that may not have previously shown significant correlations with cattle density. With the whole dataset of predictors (model I) breeding area remained the most important predictor regardless the sampling methodology used. For the other predictors of importance, they differed dramatically comparing both sampling methodologies regardless of the datasets of predictors used. This may highlight how predictor sampling methodology can influence model outcomes and the relative importance of variables. Nevertheless several studies (37–39) have questioned the relevance and possible interpretation of the PVI when correlations between predictors exist.

### Predictors

Regarding the predictors and the PCC GOF metrics (Fig.6), when using the random sampling method, there was no difference between the full predictor dataset (Model I) compared to Model II without the Karucover derived predictors. However, when using the average sampling method, an improvement in prediction accuracy was observed when using the Model I datasets. Nevertheless, as demonstrated previously, random sampling yielded superior results in both the Model I and the Model II datasets. This finding suggests the potential for achieving comparable prediction accuracy with fewer predictors. This could be particularly beneficial in the Caribbean region, where the model is intended for wider application. Even in upper-middle income countries, high-resolution land cover datasets such as Karucover are often not available, so the ability to perform well with limited predictors is a practical and valuable consideration.

## Method

In line with Da Re et al. (2020) (32), this study used the finest available census data in France—at the municipal level—for model development. However, these data are based on breeders’ declarations (40), and associate cattle with the breeders’ residence, which does not always reflect the actual location of their animals. To address this geographical bias, particularly in urban areas, the methodology employed in this study included specific adjustments. Nevertheless, the question remains as administrative boundaries do not always correspond to biological realities of cattle farming.

As highlighted by Robinson et al. (2014) (22), validating model predictions at the pixel level would require more detailed census data collected consistently over at least one year. Such an effort, however, would demand substantial coordination and resources from both the scientific community and the livestock sector.

## Conclusion

This study highlighted the value of applying census disaggregation methods to smaller territories, such as Guadeloupe, which has a relatively small number of administrative units. The maps produced by this study will have significant applications in the field of epidemiology. The high-resolution livestock density maps produced at a 225m scale represent a significant improvement over previously available GLW maps (41), which offered a resolution of 5 minute of arc (approximately 9 km x 9 km over the Guadeloupean archipelago). This work also provided an updated source of livestock data for the region, using recent census information. The methodology developed here will be adapted to other Caribbean islands; however, cattle census data and relevant predictors will need to be collected in these regions. The random sampling method of predictor selection showed advantages over the average sampling method, producing maps of comparable quality with a reduced dataset, even in the absence of high-resolution land cover data. This approach may be particularly useful in regions lacking detailed datasets such as Karucover. A potential next step to improve livestock density mapping would be to be independent of census data collected at the administrative level. Mapping densities assessments based on grazing or production areas, and explicitly identifying areas where livestock are absent such as, urban, natural or protected zones, could provide valuable insights for training datasets. This approach could also serve as a means to validate the downscaling of administrative census data to grid scale, further improving the accuracy and applicability of territorial livestock mapping models.

## Acknowledgements

We acknowledge the DAAF Guadeloupe for providing list of breeder contacts and the animal census. We would also like to thank the geological and mining research bureau (Bureau de Recherche Géologique et Minière; BRGM) of Guadeloupe for providing us with the MNT layer. This study was sponsored by the RACE project (USDA-ARS Project Number: 3022-32000-018-006-S).

## Supporting informations

**S1 Figure. Suitability raster creation processes** (Zoom on the municipality of Petit-Bourg (Prise d’eau sector, Guadeloupe).

**S2 Table. Masks used to build predictions maps and to create the suitability raster.**

**S3 Table. Symplification of level 5 Karucover 2017 land cover classes**

**S4 Table. Symplification of level 5 Karucover 2017 land use classes**

**S5 Table. Conversion of combined simplified cover & use classes to final landcover classes used for modelling**

**S6. Process of correcting the cattle density map: A) original map; B) redistribution of cattle densities from urban municipalities to neighbouring rural municipalities; C) map with superposition of the suitable areas mask; D) cattle density map with density corrected according to the surface of suitability**.

**S7. Left: Spearman correlations between cattle density and predictors; right: correlations between predictors.**

